# Emotion-related impulsivity is related to orbitofrontal cortical sulcation

**DOI:** 10.1101/2024.01.14.574481

**Authors:** William L. Hastings, Ethan H. Willbrand, Matthew V. Elliott, Sheri L. Johnson, Kevin S. Weiner

**Author notes:** **Corresponding author:** Kevin S. Weiner. **Financial Disclosures:** All authors report no biomedical financial interest or potential conflicts of interest.

## Abstract

**Background:** Emotion-related impulsivity (ERI) describes the trait-like tendency toward poor self-control when experiencing strong emotions. ERI has been shown to be elevated across psychiatric disorders and predictive of the onset and worsening of psychiatric syndromes. Recent work has correlated ERI scores with the neuroanatomy of the orbitofrontal cortex (OFC). Informed by a growing body of research indicating that the morphology of cortical folds (sulci) can produce insights into behavioral outcomes, the present study modeled the association between ERI and the sulcal morphology of OFC at a finer scale than previously conducted.

**Methods:** Analyses were conducted in a transdiagnostic sample of 118 individuals with a broad range of psychiatric syndromes. We first manually defined over 2000 sulci across the 118 participants. We then implemented a model-based LASSO regression to relate OFC sulcal morphology to ERI and test whether effects were specific to ERI as compared to non-emotion-related impulsivity.

**Results:** The LASSO regression revealed bilateral associations of ERI with the depth of eight OFC sulci. These effects were specific to ERI and were not observed in non-emotion-related impulsivity. In addition, we identified a new transverse component of the olfactory sulcus in every hemisphere that is dissociable from the longitudinal component based on anatomical features and correlation with behavior, which could serve as a new transdiagnostic biomarker.

**Conclusions:** The results of this data-driven investigation provide greater neuroanatomical and neurodevelopmental specificity on how OFC is related to ERI. As such, findings link neuroanatomical characteristics to a trait that is highly predictive of psychopathology.

## Introduction

Understanding the neuroanatomical basis of psychopathology is a major interest of cognitive and clinical neuroscience. Determining the neural correlates underlying emotion-related impulsivity (1–2) is particularly important as ERI is increasingly recognized as a shared trait across many psychological disorders (2–4) and is predictive of numerous internalizing and externalizing syndromes (5–7). Despite the well-established predictive power of ERI, neurobiological correlates of ERI are still largely unknown (8).

Previous functional and structural imaging work has suggested a link between ERI, psychopathology more broadly, and the orbitofrontal cortex (OFC). For example, previous work identified (i) a qualitative link between the pattern of cortical indentations, or sulci, in OFC and clinical outcomes (reviewed in (9)) and (ii) a quantitative link between the local gyrification of OFC and ERI (10). Nevertheless, while a growing body of work has identified quantitative relationships between sulcal morphology in different parts of the cerebral cortex with multiple behavioral and psychiatric outcomes (11–17), it is presently unknown if there is an anatomo-cognitive link between the morphology of individual sulci in OFC and ERI.

To begin to fill this gap in knowledge, we examined the relationship between ERI and OFC sulcal morphology in a transdiagnostic sample of 118 individuals with a broad range of internalizing and externalizing syndromes. To do so, we defined putative primary, secondary, and tertiary sulci in each individual hemisphere, resulting in over 2000 manually defined sulci across the sample. We then implemented a model-based LASSO regression to relate OFC sulcal morphology to ERI, and subsequently tested whether effects were specific to ERI as compared to non-emotion-related impulsivity. We focused on sulcal depth as it is one of the main morphological features differentiating putative primary (deepest), secondary, and tertiary (shallowest) sulci from one another and recent research identifies a link between sulcal depth and cognition (11–14). To our knowledge, this is the first study to examine the relationship between ERI and sulci identified in each individual hemisphere (including putative tertiary sulci, which have been largely overlooked and are considerably variable across hemispheres (15,18–19)), which is consistent with what has been referred to recently as a “precision imaging” approach (20).

## Methods

### Dataset

#### Participants

This study aims to assess the morphological characteristics of OFC sulci across a transdiagnostic sample of individuals with a wide range of internalizing and externalizing syndromes. Most participants demonstrated substantial impairment due to mental health symptoms, as indicated by receipt of disability, mental health services or by scores greater than 5 on the Sheehan Disability Scale (21); to sample a fuller range of impairment, though, 17 individuals with Sheehan Disability scores less than or equal to 5 were included.

The UC Berkeley Committee for the Protection of Human Subjects approved the parent study supported by the National Institutes of Health (NIH; Grant No. R01MH110447 [to S.L.J.]; 22) and includes 118 participants (age 18-55; 67% female, 28% male, and 5% non-binary). Researchers recruited participants through flyers, online advertising, and referrals from clinicians and excluded individuals with current alcohol or substance abuse disorders, a history of bipolar disorder or primary psychosis (as assessed by the SCID for DSM-5), or daily use of marijuana or sedating medications, as well as those with lifetime head trauma resulting in loss of consciousness for five or more minutes, low cognitive abilities (Orientation Memory Concentration Test (23) score of less than 7), MRI safety contraindications, neurological disorders, or inability to complete cognitive measures due to intellectual or language impairment. After researchers gave informed consent, participants completed diagnostic, behavioral, and neuroimaging sessions (10). Before the neuroimaging session, participants completed urine toxicology screening.

#### Behavioral Measures

The mental health-related impairment of the participants was assessed using the Sheehan Disability Scale (21). The Orientation Memory Concentration Test was used (23) to further identify any potential cognitive deficits. ERI was assessed using the well-validated Three Factor Impulsivity Index, which contains three factor-analytically derived subscales with robust internal consistency (7,24). The first factor, Feelings Trigger Action (FTA), covers the propensity to act or speak rashly while experiencing high (positively or negatively valenced) emotions. This factor is composed of three facets: the Negative Urgency Scale, Positive Urgency Measure, and Reflexive Reaction to Feelings Scale (24–26). The second factor, Pervasive Influence of Feelings (PIF), comprises three facets that assess cognitive and motivational reactions, mostly towards negative emotions: Generalization, Sadness Paralysis, and Emotions Color Worldview (24,27). The third factor, Lack of Follow Through (LFT), covers impulsivity without reference to emotion, including Lack of Perseverance and Distractibility (24,25).

The response format for all items consists of ratings on a scale of 1 “I agree a lot” to 5 “I disagree a lot.” Across previous studies, PIF and FTA, the two forms of ERI, exhibit a more substantial correlation with psychopathology measures compared to LFT (7,28,29). Given this, we hypothesized that OFC sulcal morphology would be correlated with these two factors, and we included analyses of LFT as a control measure.

### MRI Data Acquisition

Individuals participated in a brain imaging scan using a 3T Siemens TIM Trio magnetic resonance imaging (MRI) scanner with a 32-channel receiver head coil. The scanner acquired sagittal T1-weighted structural images with a standard 6.1 min. magnetization-prepared rapid gradient-echo sequence (MPRAGE) utilizing the following parameters: Repetition Time = 1900ms, Echo Time = 2.89ms, Field of View = 256mm, Voxel size = 1mm isotropic voxels, PAT Mode = GRAPPA, and PE = 2. Before the scan, researchers reminded participants to remain as still as possible, and presented participants with a blank screen during the scan (10).

### Analysis Pipeline

#### Manual definition of OFC sulci

The structural T1-weighted scans underwent processing through FreeSurfer (version 6.0.0; see https://surfer.nmr.mgh.harvard.edu; 30,31). We employed the built-in function recon-all to transform 2D high-resolution anatomical images into 3D pial and inflated cortical reconstructions. The curvature metric within FreeSurfer was used to differentiate between sulcal and gyral components (30,31,32).

W.L.H. and E.H.W. manually identified OFC sulci in each hemisphere (N = 118; 236 hemispheres; see Supplementary Figure 1) using FreeSurfer’s tksurfer tools and guided by the latest definition of OFC sulci (33). Every label was validated by a neuroanatomist (K.S.W.) prior to any morphological analyses. The OFC sulci of interest in the present study were the: (1) olfactory sulcus (olfs), (2) transverse olfactory sulcus (tolfs), (3) transverse orbital sulcus (tos), (4) anterior section of the medial orbital sulcus (mos-a), (5) posterior section of the medial orbital sulcus (mos-p), (6) anterior section of the lateral orbital sulcus (los-a), (7) posterior section of the lateral orbital sulcus (los-p), (8) intermediate orbital sulcus (ios), (9) posterior orbital sulcus (pos), and (10) sulcus fragmentosus (sf). Figure 1a provides a visual representation of left and right hemispheres for reference.

**Figure 1.**
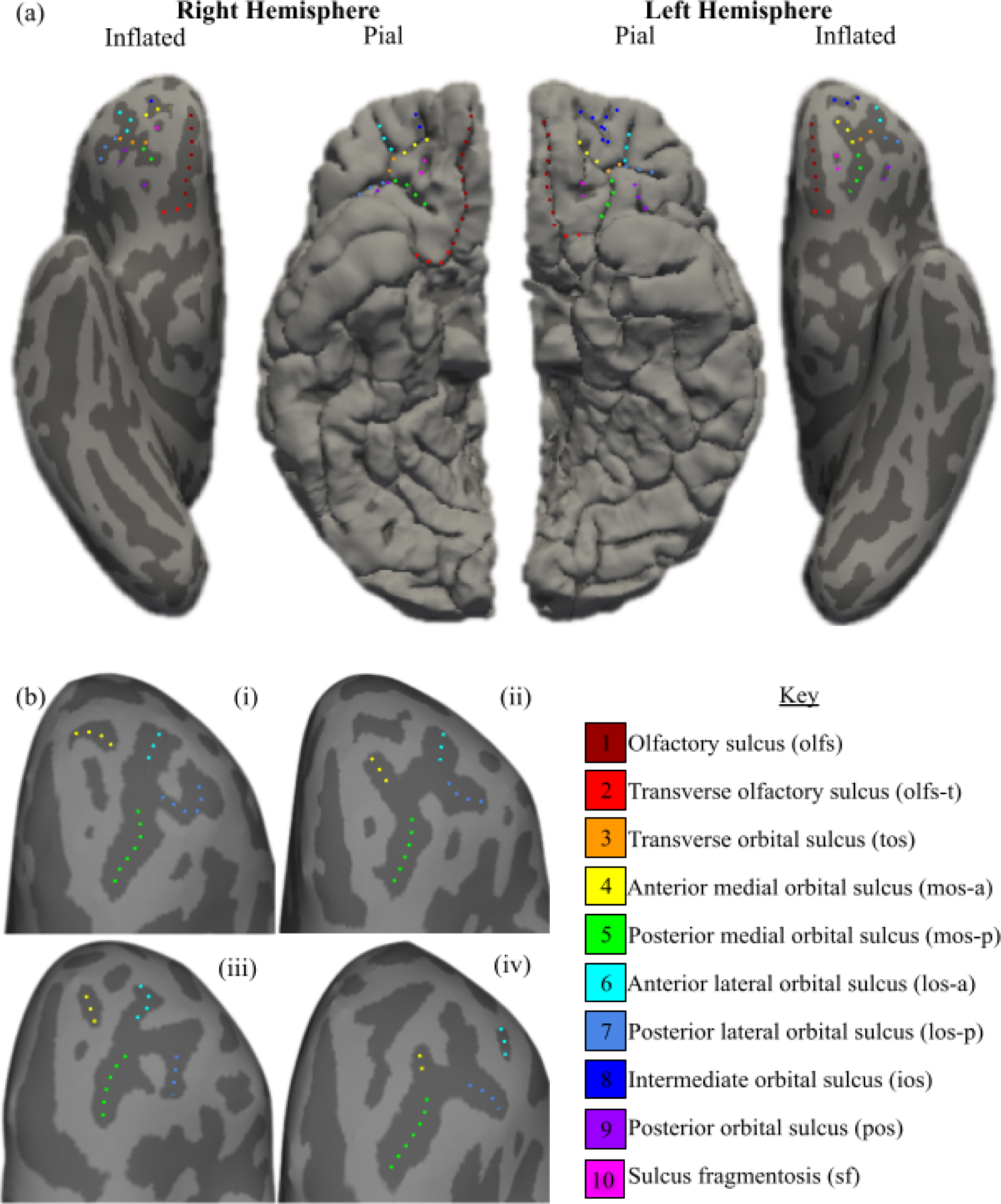
Sulci and sulcogyral types in orbitofrontal cortex. (a) OFC sulcogyral organization labeled on pial (middle surfaces) and inflated (outer surfaces) surfaces in the left (right surfaces) and right (left surfaces) hemispheres. On these cortical reconstructions, sulci are dark gray and gyri are light gray. Sulci are outlined according to the key beside b. Sulci are labeled according to a revised version of Petrides’ (33) atlas. Example hemispheres shown have all variable sulci, which is not always the case with participants in this study. (b) Examples of the different OFC patterns as defined by Chiavaras and Petrides (34) and refined by Chakirova and colleagues (35).

#### Characterizing OFC sulcogyral patterns into “Types”

We categorized each hemisphere’s sulcogyral organization based on a classification system developed by Chiavaras and Petrides (34) and refined by Chakirova and colleagues (35) (Figure 1b i-iv). Type I, the most prevalent sulcal pattern across samples (9), displays a discontinuity between the anterior and posterior components of the medial orbital sulci, while the anterior and posterior components of the lateral orbital sulcus are continuous. Type II demonstrates continuity between the anterior and posterior components in both the medial and lateral orbital sulci. Type III shows discontinuity between the anterior and posterior components of both the medial and lateral orbital sulci. Type IV, the least common and typically excluded from analysis, exhibits discontinuity in the anterior and posterior components of the lateral orbital sulci, but not the medial orbital sulci. We also assessed the number of variable putative tertiary sulci — sulci with an incidence rate that varies from zero to up to four components across individuals — in each hemisphere. Similar to the labeling process, each pattern underwent assessment and validation before any further analyses were performed.

#### Extracting Morphological Features

After sulci were defined, depth (according to FreeSurfer values (30), cortical thickness (in millimeters), and surface area (in square millimeters; 31) were calculated using an established analysis pipeline (12,20) and the mris_anatomical_stats function in FreeSurfer (32). These metrics commonly discriminate sulcal types from one another. For example, primary sulci are the deepest with the largest surface area, while putative tertiary sulci are the shallowest with the smallest surface area (36–38). The latter are particularly intriguing when considering neurodevelopmental disorders as they emerge last in gestation in association cortices and continue to develop after birth. Depth values, as computed by FreeSurfer, depend on the distance of a vertex from the “mid-surface,” with the mean displacements around this “mid-surface” being zero. This usually results in gyri having negative values, while sulci have positive values. However, due to the shallowness and variability in the depth of putative tertiary sulci, some mean depth values can be less than zero. To account for variations in brain size among individuals and hemispheres, we employed normalized sulcal depth, consistent with previous studies (11,12).

### Data Analysis

#### Continuous Measurements

Building upon our prior work (11,12,13), to analyze continuous morphological sulcal-behavior relationships, we leveraged a pre-existing pipeline that tests for a relationship between sulcal morphology (i.e., sulcal depth) and ERI via the following approach:

1. L1 regularization (LASSO regression): this data-driven method selects a model which filters out regressors that contribute below a threshold predictive value, safeguarding against overfitting and maximizing generalizability. The result is a parsimonious, yet comprehensive, model.
2. Cross-validation: We then subject the resulting model to two stages of cross-validation.

First, we determine the optimum shrinking parameter (alpha value) based on what value minimizes the mean squared error (MSE) of the model. Next, we fit the models using standard leave-one-participant-out cross-validation to obtain cross-validated R2 and MSE values. Finally, to further assess the model we obtained bootstrapped median and 95% confidence intervals for MSE.

This approach requires that all sulci are present in each hemisphere. As such, to balance the number of sulci and number of participants in the models, we excluded the pos and ios as they were the most variable (Figure S2). In total, morphological data from 109 left hemispheres and 102 right hemispheres were included in the models.

We tested for a relationship between OFC sulcal morphology with the two emotionally driven ERI factors (PIF and FTA) in each hemisphere separately. The data-driven LASSO regression selected a subset of ten sulci in each hemisphere based on the predictive value of their depth for FTA. The models were then compared to two alternative models to assess specificity. The first model acted as a surface-based measurement control (cortical thickness) and the second served as a behavioral control (Lack of Follow Through). We assessed how our model performed relative to the control models by measuring the difference of their Akaike Information Criterion (AIC; 39).

#### Categorical Measurements

In addition to the model-based approach described in the previous section, we also considered the relationship between ERI and two categorical measures. The first was the Type of OFC sulcogyral pattern. The second was the total number of variable (putative tertiary) OFC sulci. For these two categorical measures, we conducted a Type III Analysis of Variance (ANOVA) to compare incidence rates for each ERI factor. We used Post-hoc Tukey tests to test for differences between group means.

## Results

### Overview of Orbitofrontal Sulcal Incidence and Sulcogyral Patterning

We were able to identify 9 sulci that have been previously identified and explored, six of which were identifiable in every hemisphere: 1) the posterior and anterior portions of the lateral (los-a, los-p) orbital sulcus, 2) the anterior and posterior portions of the medial (mos-a, mos-p) orbital sulcus, 3) the transverse orbital sulcus (tos), and 4) the olfactory sulcus (olfs). Three additional sulci could be defined in some, but not all, hemispheres across participants: (1) the intermediate orbital sulcus (ios), (2) the posterior orbital sulcus (pos), and (3) sulcus fragmentosis (sf). Additionally, these sulci varied in the number of observed components, ranging from zero to as many as four components (Figures. 1a, 5b-c).

For the first time (to our knowledge), we were also able to identify the transverse olfactory sulcus (Figure 1a; tolfs) in every hemisphere examined. Though many neuroanatomists throughout history have acknowledged a “hook” at the posterior extent of the olfactory sulcus (Supplementary Materials; Supplementary Figures 5-6), the present study is the first (to our knowledge) to define and label this component separately. Comparing cortical thickness and depth between the olfs and tolfs revealed that the latter is significantly deeper and thicker (Tukey p < 0.0001 for all comparisons) than the former (Figure 2), providing empirical evidence that the tolfs is morphologically distinct from the olfs.

**Figure 2.**
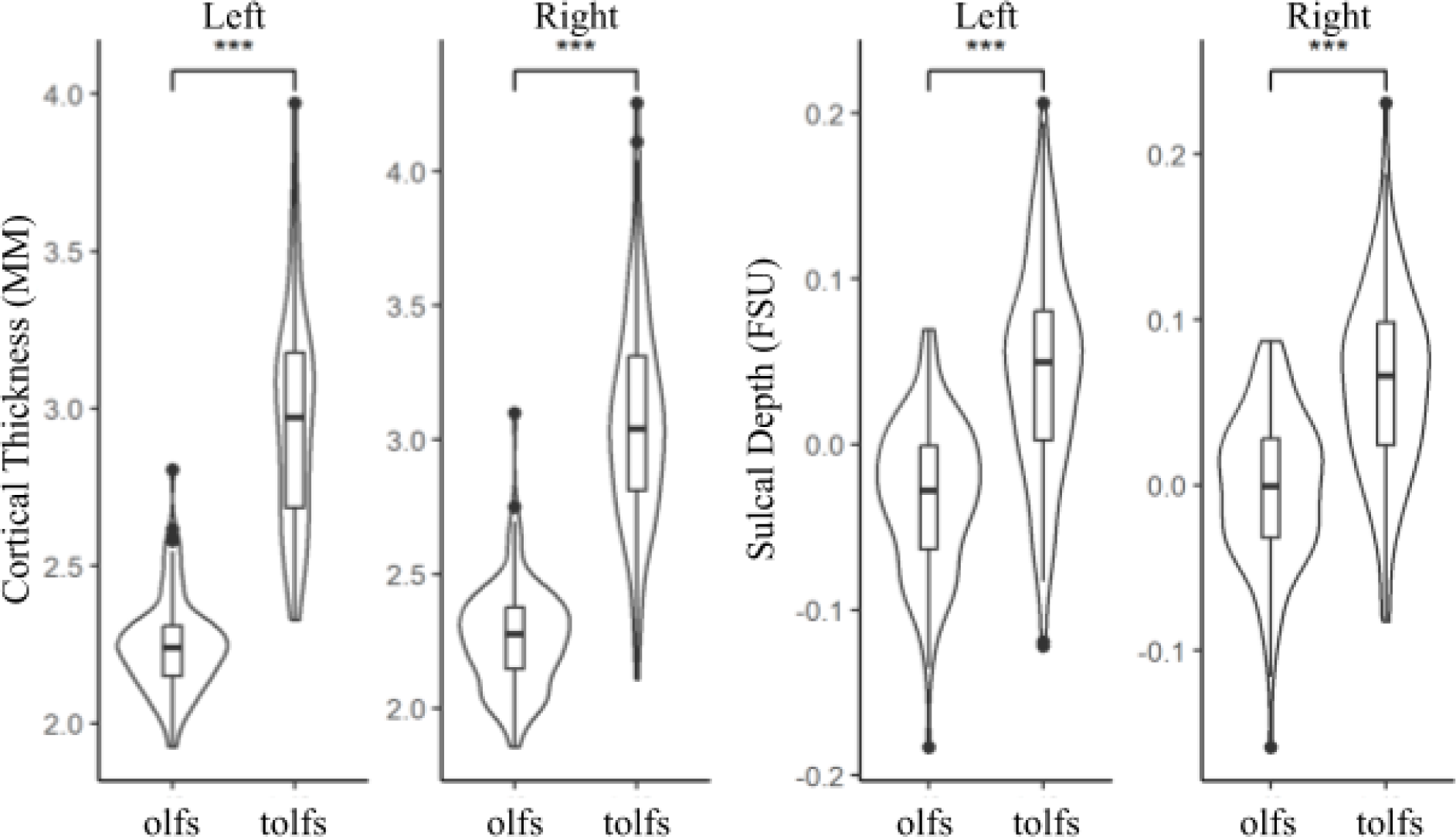
The transverse olfactory sulcus is cortically thicker and deeper than the olfactory sulcus. Distributions are represented by box plots. Outliers are represented as points. Significant differences are denoted by bars above the graph. Cortical thickness is measured in millimeters (mm) and sulcal depth is measured in FreeSurfer Units (FSU; see materials and methods for additional details). Sulcal abbreviations correspond to those used in Figure 1. *** p < 0.0001

OFC sulcogyral patterns also varied (Figs. 1b, 5a) with comparable incidence rates reported first by Chiavaras and Petrides (34) and by subsequent studies (see 9 for review). To directly compare our incidence rates to those previously reported, we performed a meta-analysis comparing the incidence rates in the present study to studies reporting OFC sulcogyral patterns in neurotypical controls (12 studies, 710 total participants), and clinical samples (13 studies, 869 total participants). The prior clinical samples comprised individuals with first episode psychosis, ultra-high risk of psychosis, schizophrenia, schizotypal, and autism spectrum disorder, all of which have reported sulcogryal incidence rates (9,34–35,40–49) (Figure 3). Finally, ANOVAs examining the effect of sulcogyral type (types I-III; Figure 1; Supplementary Figure 3a) and number of variable sulci (pos, ios, and sf; Figure 1; Supplementary Figure 3b-d) on each factor of ERI revealed no significant effects across all comparisons (ps > 0.05).

**Figure 3.**
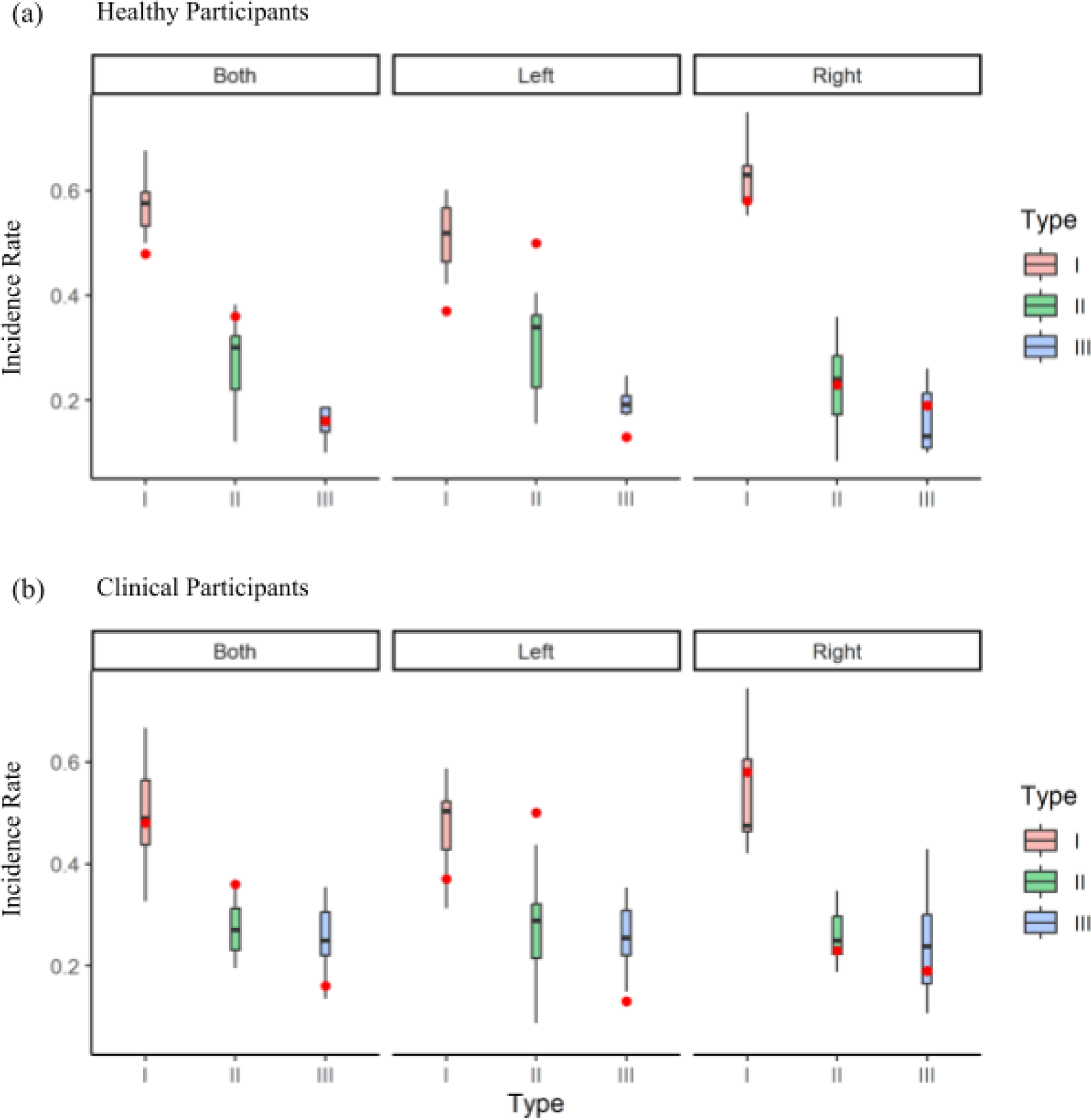
OFC sulcogyral types in the present study are consistent with a previous meta-analysis. Red points represent the incidence rate observed in this sample in comparison to boxplots of incidence rates observed across several studies in both hemispheres as well as left and right hemispheres individually in (a) healthy (12 studies, 710 total participants), and (b) clinical samples (13 studies, 869 total participants). Type IV was excluded due to inconsistent inclusion across the literature.

### The depths of a subset of OFC sulci significantly correlates with ERI

We implemented data-driven LASSO regression models using the sulcal depths of OFC sulci (tos, tolfs, sf, olfs, mosp, mosa, losp, losa) in the left and right hemispheres separately to predict each emotionally related ERI factor: (i) Feelings Trigger Action (FTA) and (ii) Pervasive Influence of Feelings (PIF). From these models, the OFC sulcal depths in the left and right hemispheres independently related with FTA, but not PIF. More specifically, the depths of eight OFC sulci between the two hemispheres correlated with FTA at the optimum alpha threshold of the model (left hemisphere: α = 0.005, MSE = 0.53; right hemisphere: α = 0.005, MSE = 0.51; Figure 4). In the right hemisphere, the depth of the olfactory sulcus (β = 0.23), as well as the posterior component of both the lateral (β = 1.82) and medial (β = −0.80) orbital sulci correlated with FTA (R2cv = 0.05, MSEcv = 0.50; Figure 4, top left). In the left hemisphere, the depth of the sulcus fragmentosis (β = −1.38), olfactory sulcus (β = −1.03), anterior (β = 0.90) and posterior (β = 0.42) components of the medial orbital sulcus, as well as the anterior component of the lateral orbital sulcus (β = −0.34) correlated with FTA (R2cv = 0.07, MSEcv = 0.49; Figure 4, bottom left). In both hemispheres, we observed a modest correspondence between predicted and actual measured FTA scores (left hemisphere: Spearman’s rho = 0.26; right hemisphere: Spearman’s rho = 0.24).

**Figure 4.**
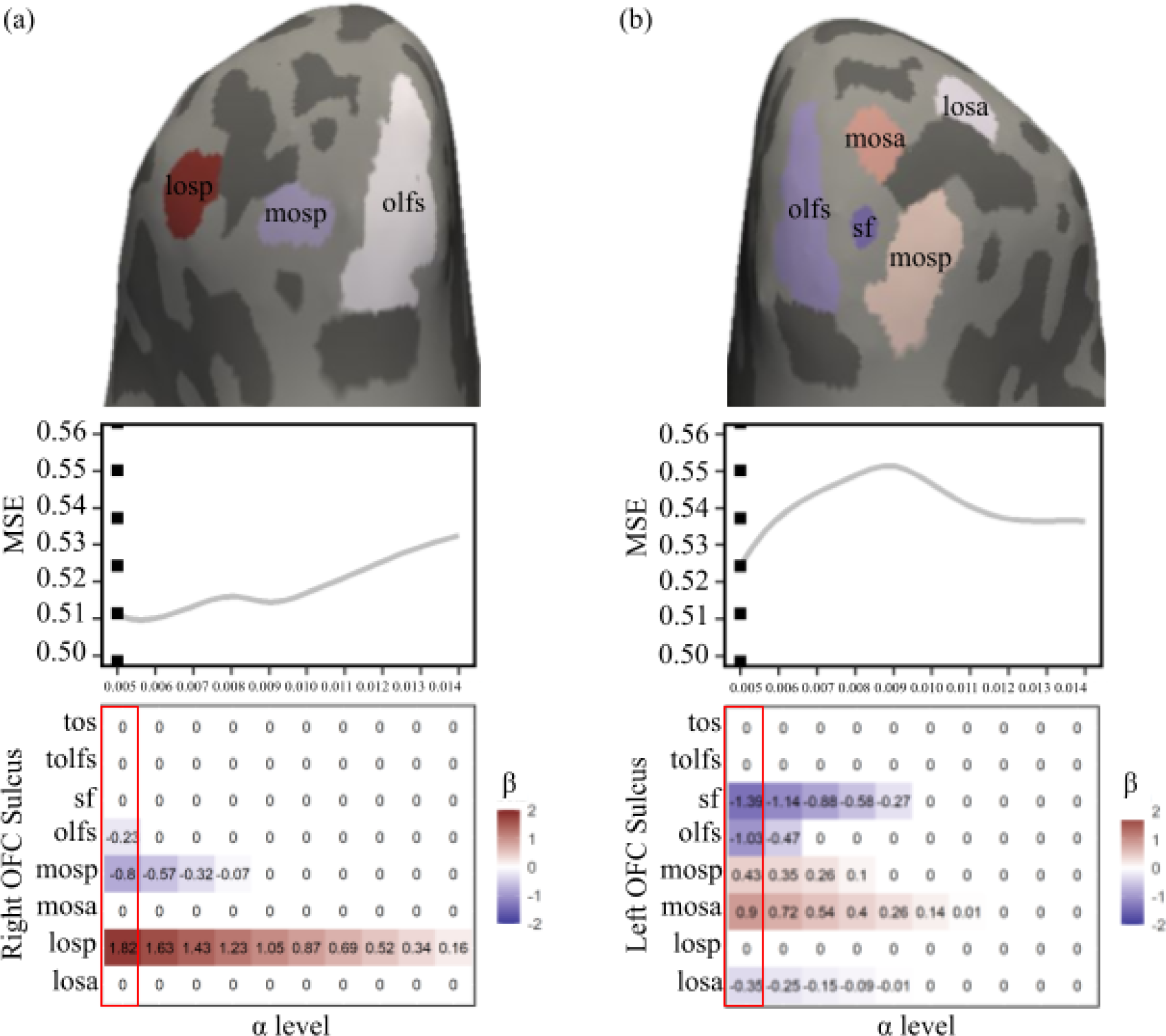
Data-driven model selection reveals a relationship between OFC sulcal depth and emotion-related impulsivity. Inflated cortical surfaces including LASSO selected sulci (top) colored according to the beta values (bottom). Line graphs depict MSE at each corresponding alpha (middle). The dotted line represents the alpha which minimizes MSE. Matrices reflect the beta values of the predictors at each corresponding alpha value (bottom). Beta values used in the model which minimize MSE are outlined in red. Sulcal abbreviations correspond to those used in Figure 1.

The Akaike Information Criterion (39) assesses how well the depth models predict FTA in comparison to our control measures for both behavior and morphology. If the ΔAIC is greater than 2, it suggests an interpretable difference between models (50–51). The difference between the behavioral control measure, Lack of Follow Through (LFT), and depth (left hemisphere: △AICFTA-LFT = 7.18; right hemisphere: △AICFTA-LFT = 6.40) indicates that the predictive value of the FTA is significantly stronger than its non-emotional counterpart, LFT. Such is also the case with regard to the structural control measure and cortical thickness (left hemisphere: △AICDepth-Thickness = 5.78; right hemisphere: △AICDepth-Thickness = 2.71).

Finally, we tested the impact of defining the tolfs separately from the olfs on our models. To do so, we generated a separate model which did not differentiate the transverse portion from the main branch of the olfs. This “olfactory complex” had no predictive value on the model in the right hemisphere and decreased the beta value of the olfs in the left hemisphere by 98% (Supplementary Figure 3). This provides additional empirical evidence that the tolfs should be defined separately from the olfs.

## Discussion

In the present study, we examined the relationship between OFC sulcal morphology and ERI in a transdiagnostic clinical sample. To do so, we manually defined over 2,000 sulci in individual participants and implemented a model-based approach, which revealed bilateral associations of ERI with the depths of eight OFC sulci. These effects were specific to ERI and were not observed in non-emotion-related impulsivity. In addition, we identified a new transverse component of the olfactory sulcus in every hemisphere that is dissociable from the longitudinal component based on anatomical features and correlation with behavior. The results of this data-driven investigation provide greater neuroanatomical and neurodevelopmental specificity on how OFC is related to ERI than previous studies, as well as link neuroanatomical characteristics to a trait that is highly predictive of psychopathology. Together, our findings provide an important step in clarifying the neuroanatomical correlates of ERI, a foundation that can be built upon in future studies. In the sections below, we discuss these results in the context of previous findings, as well as consider goals for future work.

In previous work, Elliott and colleagues established a neuroanatomical link between OFC and ERI (10). Using a continuous global surface-based morphological measurement, the authors identified a link between ERI and the local gyrification of OFC. Building on that work which used an automated algorithm to define the perimeter of a large OFC region of interest (ROI), here, we improved the spatial scale of our measurements from one large OFC ROI to manually defined OFC sulci in each individual hemisphere (in line with what has been referred to as a “precision imaging” approach; 20). As such, the present findings identify precise neuroanatomical sulcal landmarks that are related to ERI and complement the previous findings. For example, Elliott and colleagues identified a stronger relationship between FTA and the OFC in the left hemisphere. Our model-based approach also identified a hemispheric difference in which the depths of 5 sulci in the left hemisphere and only 3 in the right hemisphere were related to ERI that provide two key insights. First, the model identified only two sulci (olfs and mos-p) in both the right and left hemispheres, indicating bilateral sulcal landmarks related to ERI for the first time. To remind the reader, it is important to note that this olfs definition does not include the tolfs; when including the tolfs in the definition of an “olfs complex,” the model fit plummets (see Supplementary Materials for further discussion regarding the tolfs; Supplementary Figures 4-6). As prior findings show that the depth of the olfs is typically decreased in patients with or at risk of developing schizophrenia (52–55),, our findings indicate that the olfs, but not the tolfs, could be driving those effects, which can be examined in future research. Second, the model identifies both large, primary sulci and small, putative tertiary sulci such as the sulcus fragmentosus (sf). Thus, it is not just primary, secondary, or tertiary sulci that are driving the effects previously identified; instead, it is a sulcal landscape of eight OFC sulci between the two hemispheres that are related to ERI.

There are also differences between the present and previous findings (10) indicating that the neuroanatomical correlates of FTA and PIF may operate on different spatial scales. For example, there was no significant relationship of PIF with sulcal depth, whereas Elliott and colleagues (10) found that PIF correlated bilaterally to the local gyrification of OFC. As such, PIF may function at a coarser level in OFC (broad, non-sulcal-specific folding) while FTA appears to be localized to specific structures. In a similar vein, and more broadly, our present findings also indicate that ERI as a construct relates more to fine-grained, continuous features of the cortex that coarser, categorical measures like sulcal incidence can not properly capture. For example, the incidence rate of variable sulci was not significantly related to impulsivity levels, despite prior work indicating that the number of variable sulci present in clinical populations relates to psychopathology severity (for review, 9). Taken together, the combination of the present and previous findings underscore how psychopathology manifests in the OFC on multiple scales, some of which appear to overlap with impulsivity — as observed in global and local continuous measurement scales with respect to ERI — while others do not.

By focusing on sulci within OFC, we have improved the spatial scale of the neuroanatomical underpinnings of ERI. Nevertheless, it may be tempting to also conclude that these structures are the most important neuroanatomical link to ERI (i.e., a more localized “modular” view). We emphasize that the neuroanatomical underpinnings of ERI likely include many more neuroanatomical structures across spatial scales and that our present findings are the next step in uncovering the infrastructure of a complex neural network underlying ERI that is presently unknown (56). Indeed, previous findings relating sulcal morphology to cognition in different populations discuss the relationship between sulcal morphology and network connectivity with an emphasis on white matter architecture (11,12,15,57–60). The present work identifies a local sulcal network in OFC bilaterally related to ERI that will serve as the foundation for identifying additional components of the complex neuroanatomical network underlying ERI for decades to come.

Zooming out, the present results appear to further bridge parallel transdiagnostic literatures in psychiatry and neuroscience. For example, while ERI has been well-explored and accepted as a transdiagnostic phenotype in clinical literature (61–63), explorations of OFC anatomy and psychopathology appears to be less explored, with few studies testing for similarities across diagnostic boundaries (64–68). To bridge between these parallel tracks, we recently proposed that ERI could serve as an intermediate psychological phenotype that emerges from the development of OFC and leads to psychopathology (10). Here, we extend this proposal to also include the emergence of neuroanatomical structures. That is, sulci emerge at different time points in gestation and a classic theory proposes that in a given cortical expanse, sulci that emerge later will be related to aspects of cognition that have a protracted development (69–70). Consistent with this idea, Chi and colleagues (36) observed that the posterior extent of the olfs — consistent with the location of the tolfs in the present study — emerged first (around 16 gestational weeks), while the more anterior longitudinal component emerged significantly later (around 25 gestational weeks; see Supplementary Materials). The fact that our model identified the olfs, but not the tolfs, bilaterally suggests the intriguing possibility that the later emergence and development of the olfs, not the tolfs, could be related to ERI. Longitudinal research will be needed to test this hypothesis. We are hopeful that both the neuroanatomical and model-based approach and the empirical results from this study can support future work on ERI and generalize to other research domains that seek to integrate the transdiagnostic study of psychopathology at two crucial levels of analysis, psychiatry, and neuroscience. To expedite this goal, tools are actively being developed to leverage deep learning algorithms as a guide to semi-automate the definition of neuroanatomical structures such as the small and variable tertiary sulci explored here (71,72). Such tools will be sure to increase the sample size, which is a main limitation of the present study, as well as shed further light on the relationship between tertiary sulci and psychopathology. That is, despite Sanides’ classic hypothesis (69–70), tertiary sulci have been widely overlooked for methodological reasons (18 for review) and the broad fields of psychiatry and neuroscience know very little about their development, morphological changes across the lifespan, and if/how the development and changes across the lifespan relate to psychiatric outcomes. An important next step will be to include tertiary sulci in neuroanatomical investigations in different clinical populations, which could have implications for mental health treatment. For example, while one of the smallest and shallowest sulci in OFC, the depth of the sulcus framentosus (sf; Figure 4; Supplementary Figure 2b) explained the most variance of all left hemisphere sulci identified by our model. Lastly, considering that the structure and function of the prefrontal cortex (PFC) is related to impulsivity in patients with schizophrenia (73) and contains numerous tertiary sulci (11–13,33,60,74), future work should also assess the relationship between PFC sulcal morphology and ERI.

Finally, as OFC and ERI have both been responsive to existing treatments spanning cognitive behavioral therapy (8,75), cognitive training (76–78), and mindfulness (79–81), the present findings suggest that future intervention studies targeting the sulci identified here may be promising for treating psychopathology transdiagnostically.

## Acknowledgments

We thank W. Voorhies for developing the data analysis pipeline implemented in the present work, as well as K. Timpano, K. Modavi, A. Dev, M. Robison, J. Mostajabi, S. Esmail, and B. Weinberg for their help with recruitment and data collection. We also thank N. Angelides, H.Y. Tsai, M. Andrews, and J. Giffin for their help with magnetic resonance imaging data acquisition.

## Financial Disclosures

This work was supported by the National Institutes of Health (Grant No. R01 MH110447 [to SLJ]), the Brain and Behavior Research Foundation (NARSAD 30738 to KSW), National Science Foundation (NSF CAREER 2042251 to KSW), and the National Institutes of Health Medical Scientist Training Program Grant (T32 GM140935 to EHW). The funding agencies did not have a role in the study design, data collection and analysis, decision to publish, or preparation of the manuscript.

### Supplementary Methods and Materials - The Transverse Olfactory Sulcus

The olfactory sulcus has historically been described as the sulcus that “occupies the most medial position [of the orbitofrontal cortex] and runs in a straight rostrocaudal direction underneath the olfactory tract”(34). Nevertheless, over the past century, prior work has qualitatively described the posterior extent of the olfactory sulcus — for example, as “hook-like”(34,82–84). Clarifying these qualitative descriptions, here, we quantitatively showed that the posterior, transverse extent of the olfs is morphologically distinct from the anterior, longitudinal component based on thickness and depth (Figure 2). Furthermore, defining the tolfs separately impacted our models relating OFC sulcal depths to ERI (Supplementary Figure 3). As such, we define it separately as the “transverse olfactory sulcus” (Figure 1).

An additional reason to define the tolfs separately is that it putatively emerges first before the main body of the olfs. Specifically, Chi and colleagues (36) observed that the posterior extent of the olfs emerges first (at around 16 gestational weeks) and then develops from back to front (becoming prominent by gestational week 25). This would suggest that the tolfs actually emerges first, which, given Sanides’(70) theory proposing that later emerging sulci in association cortices are related to later developing aspects of cognition, could explain why our model pulls out the body of the olfs (which emerges later), but not the tolfs (which emerges first). Furthermore, previous research specifies that the posterior extent of the olfs identifies the “olfactory trigone” — a subcortical divergence of the olfactory tract into lateral, intermediate, and medial striae each of which sends signals to other cortical locations relating the sense of smell throughout the brain.

**Supplementary Figure 1.**
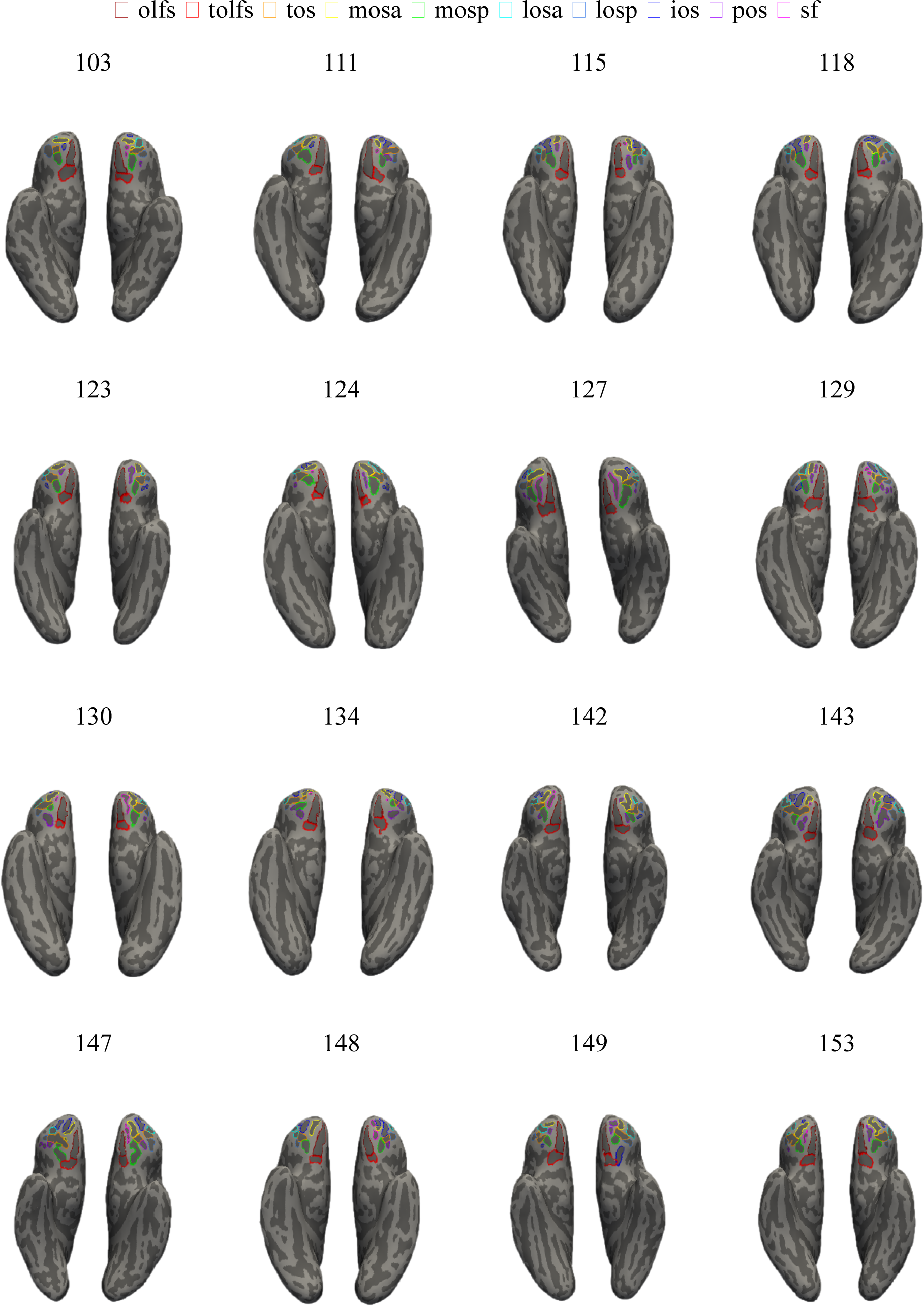
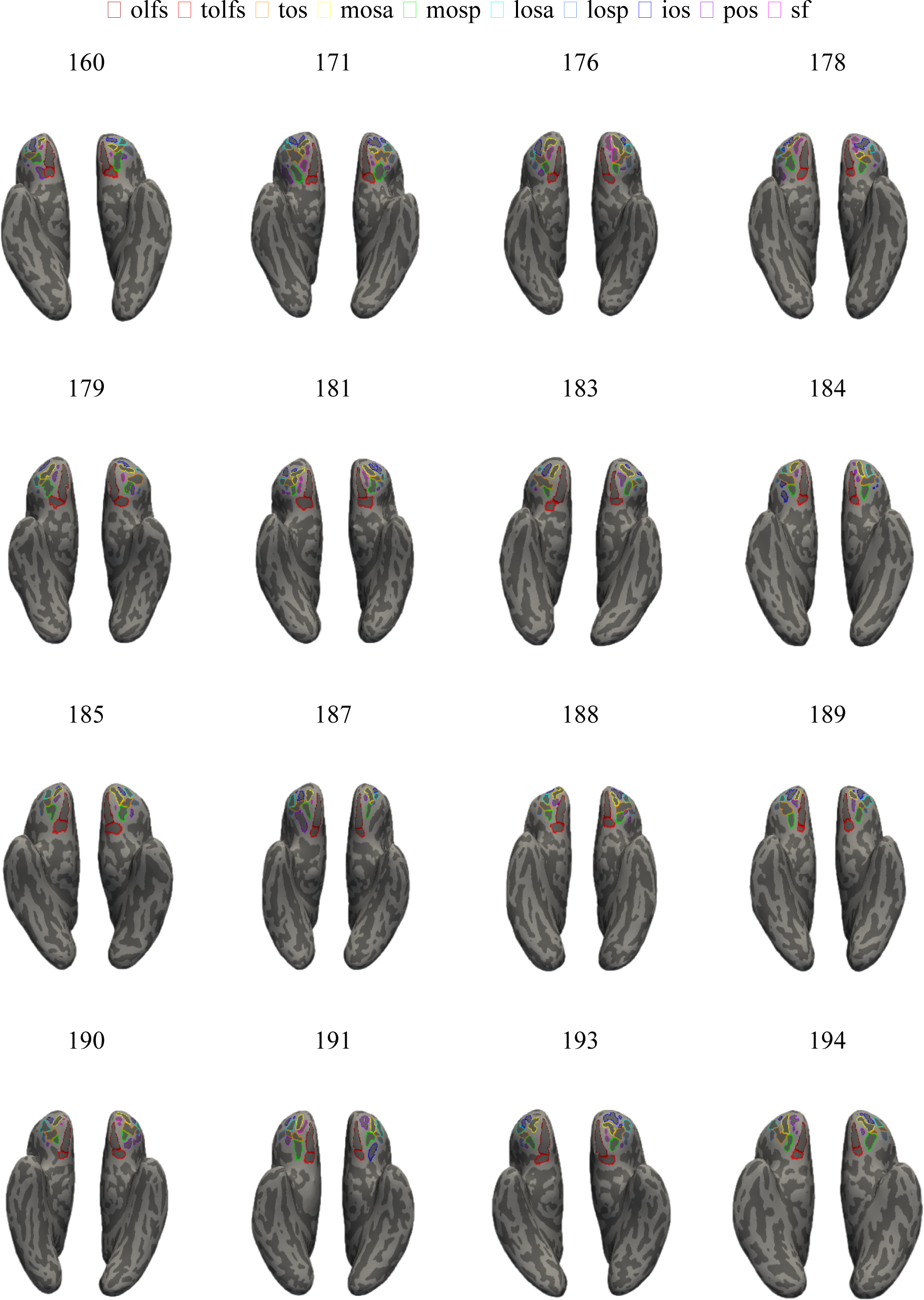
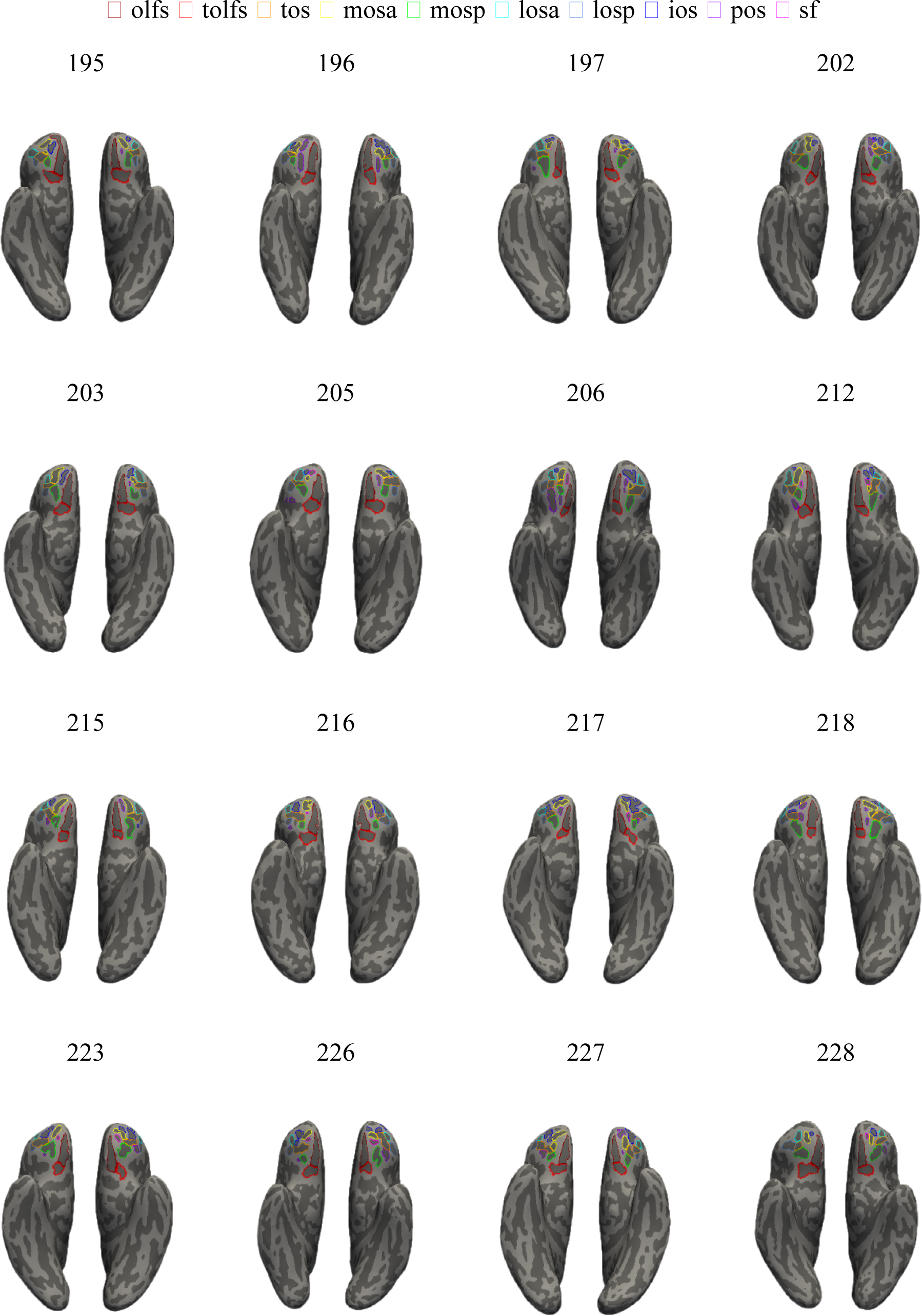
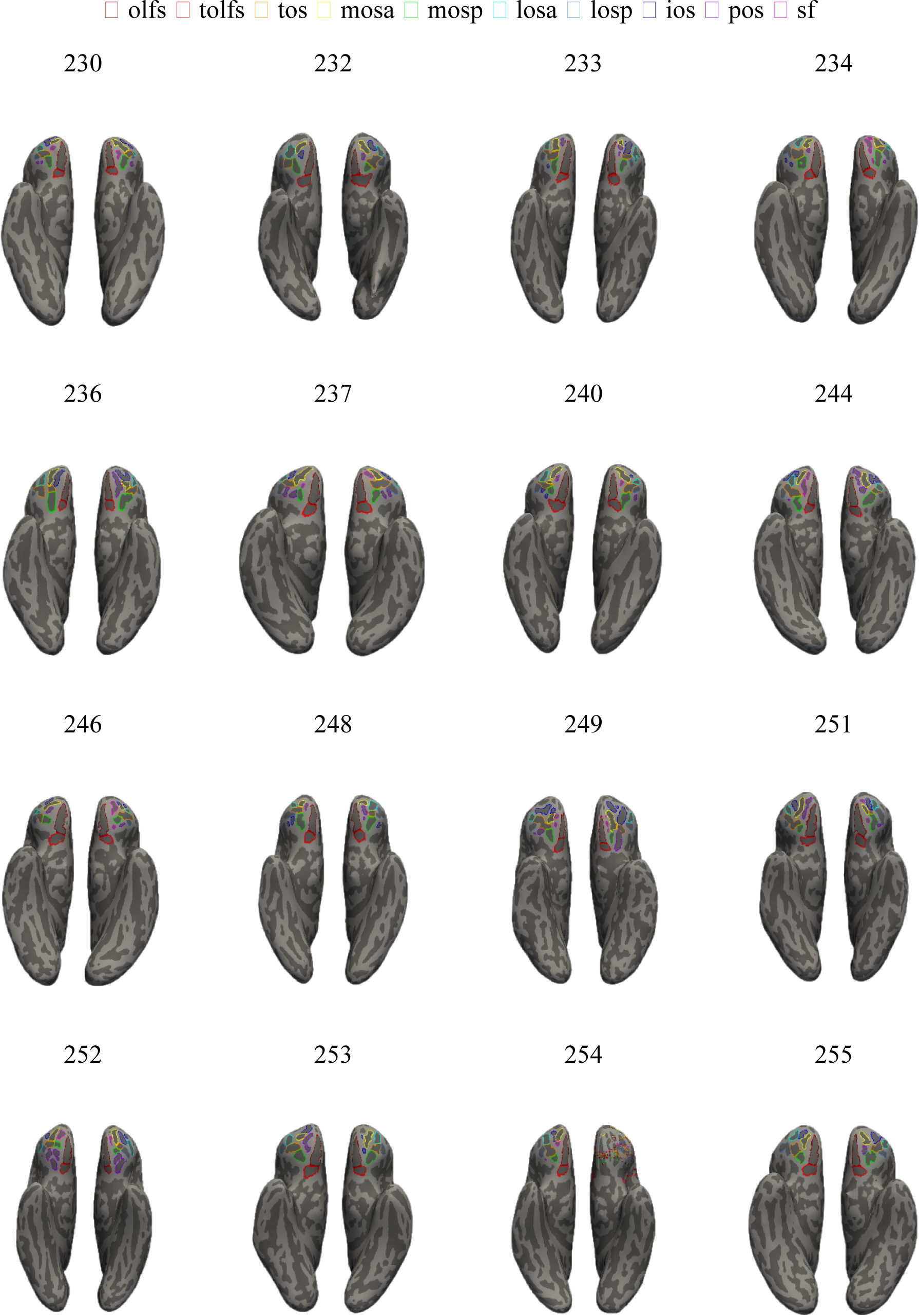
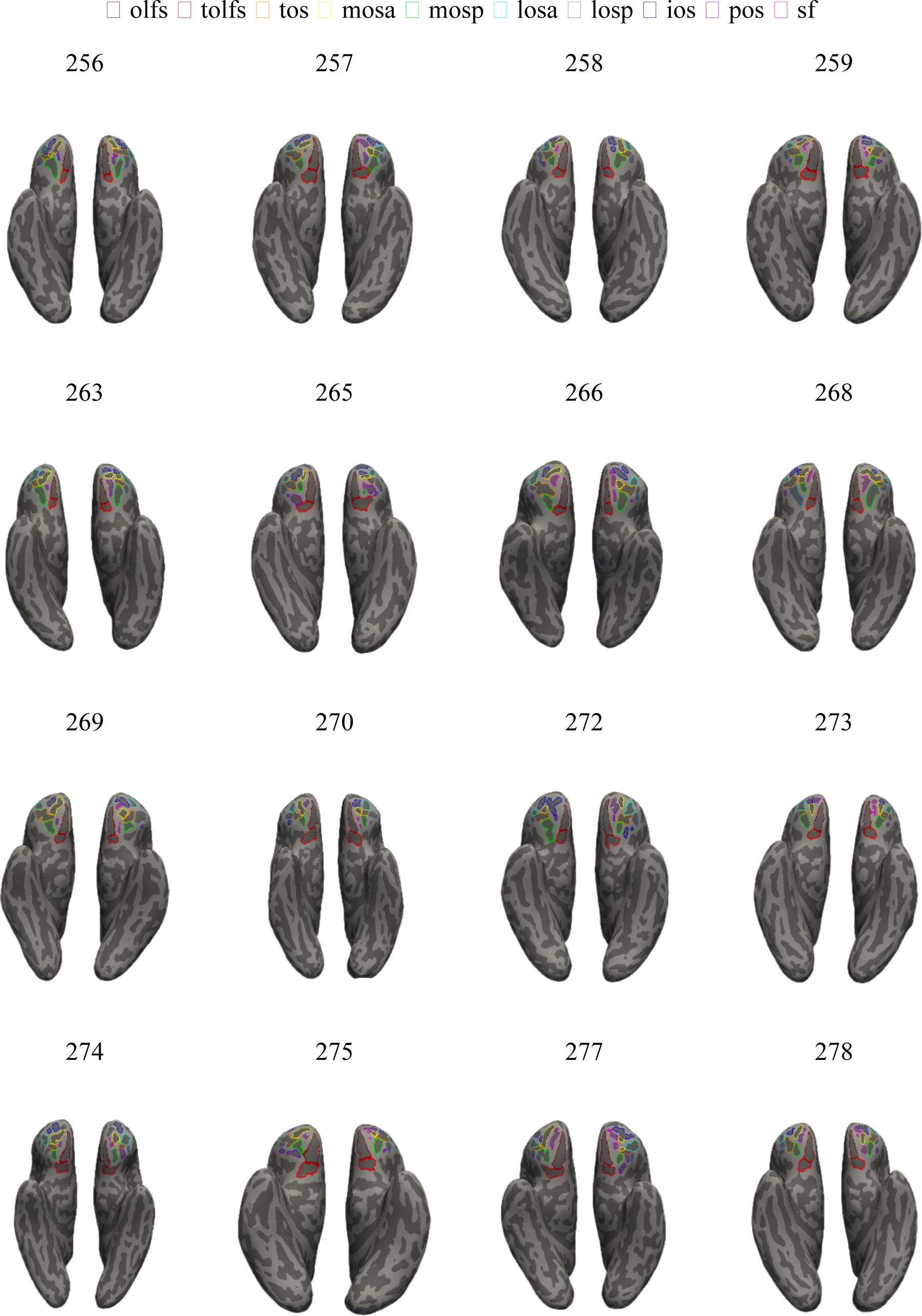
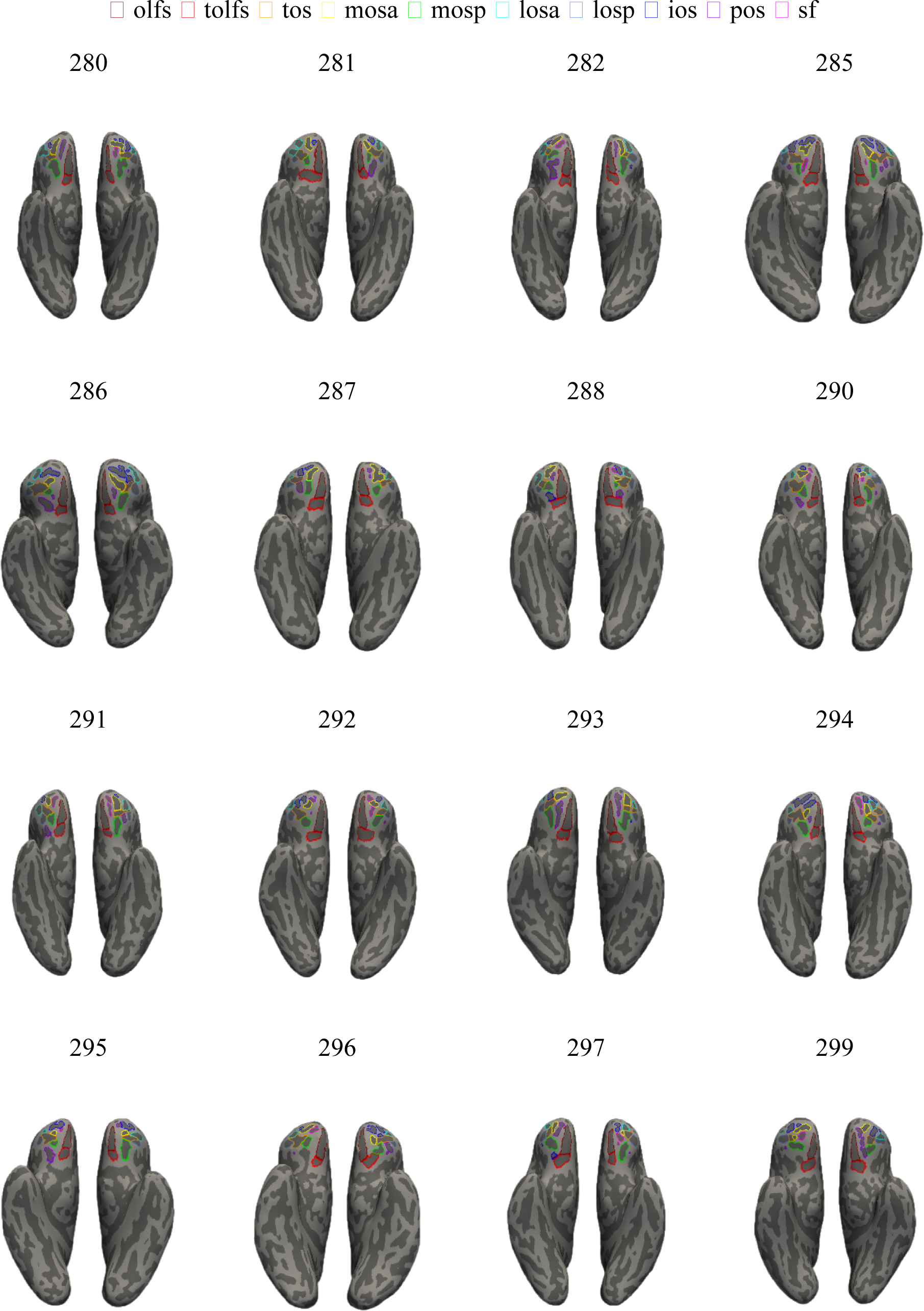
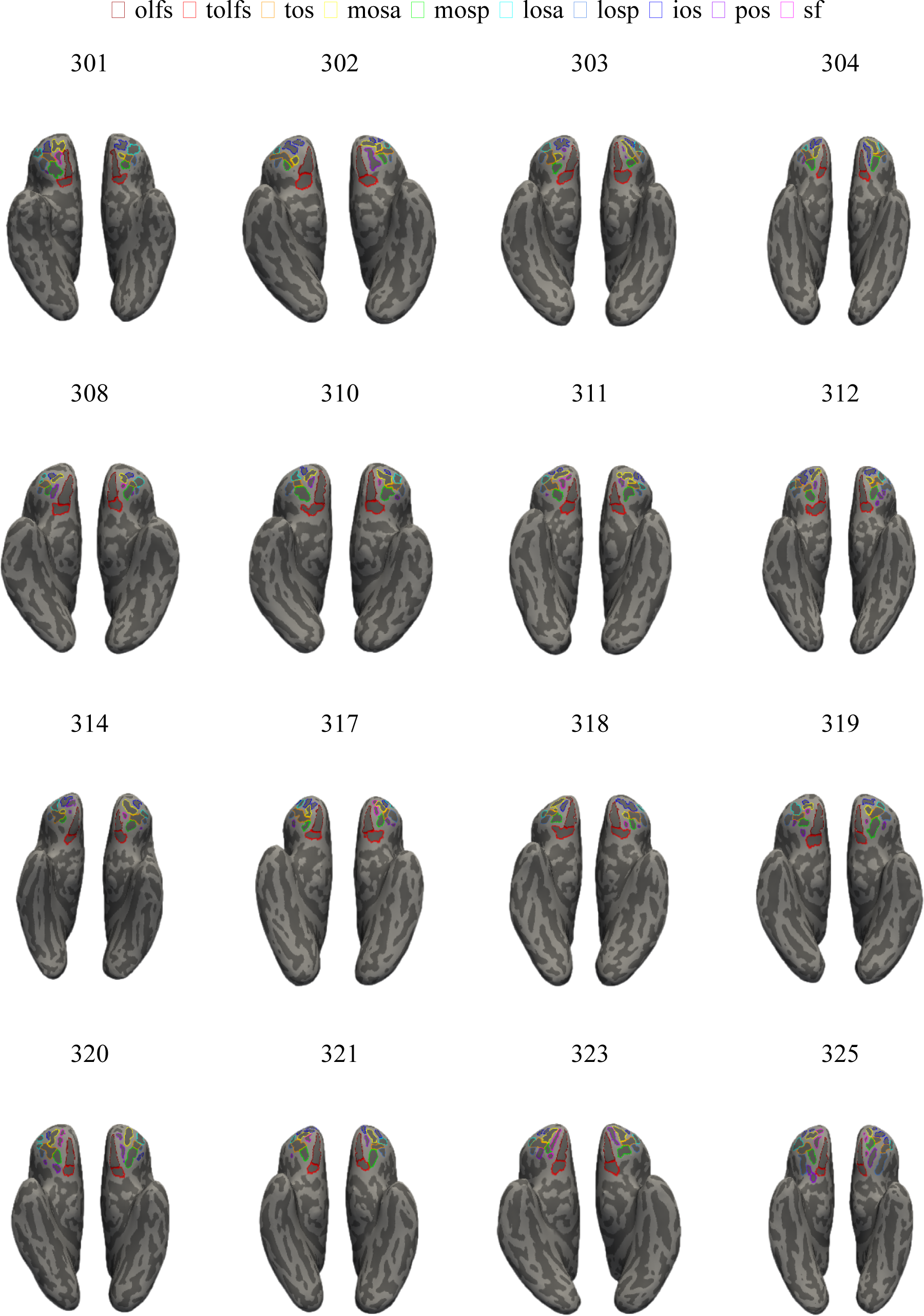
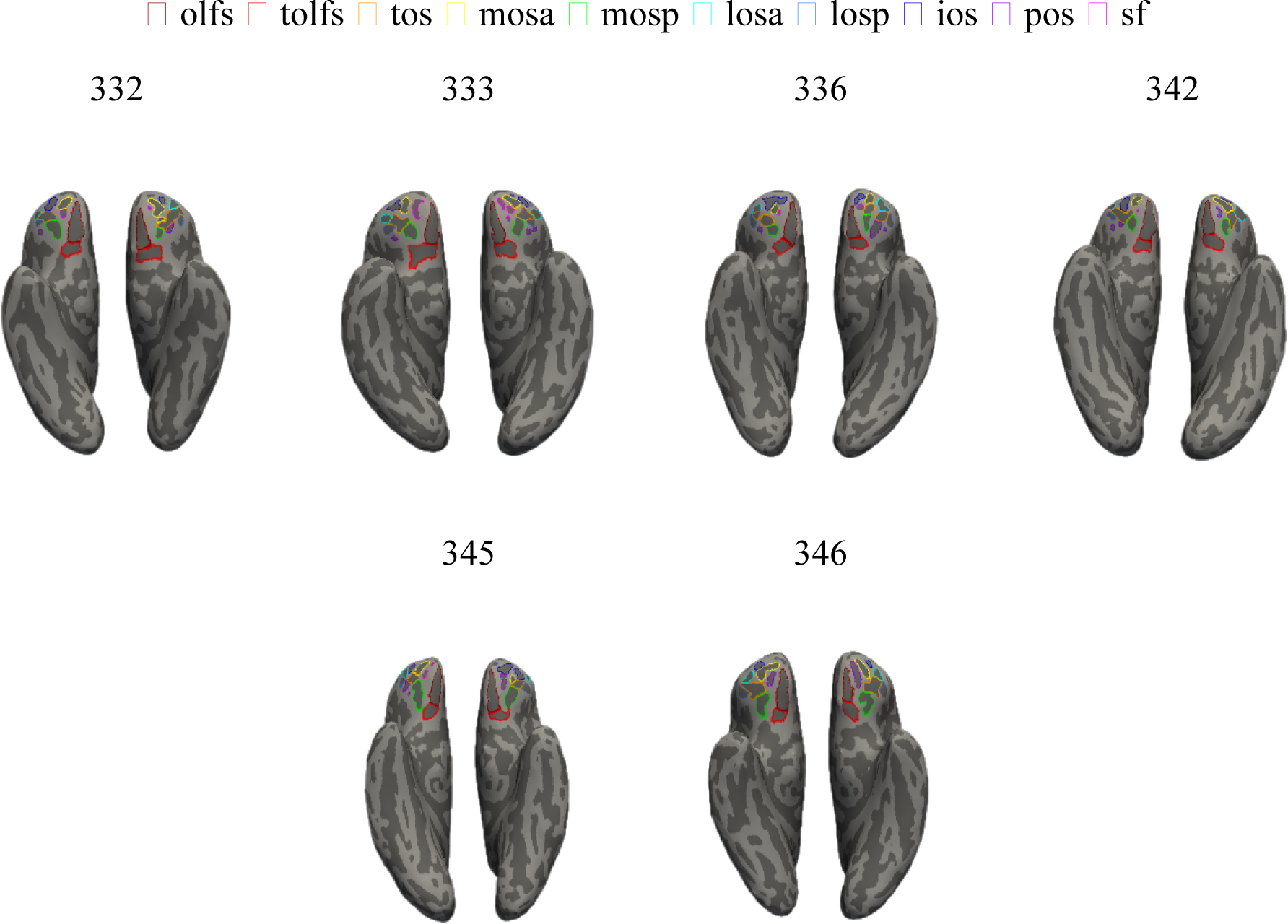
Manually defined OFC sulci in every hemisphere. Each sulcus is displayed on the left and right hemisphere inflated cortical surfaces in FreeSurfer 6.0.0, with labels displayed as an outline according to the key at the top. Each hemisphere contains at least 7 sulci (from medial to lateral, anterior to posterior): olfs, tolfs, mosa, mosp, tos, losa, and losp. An additional 3 sulci are variably present: sf, ios, pos.

**Supplementary Figure 2.**
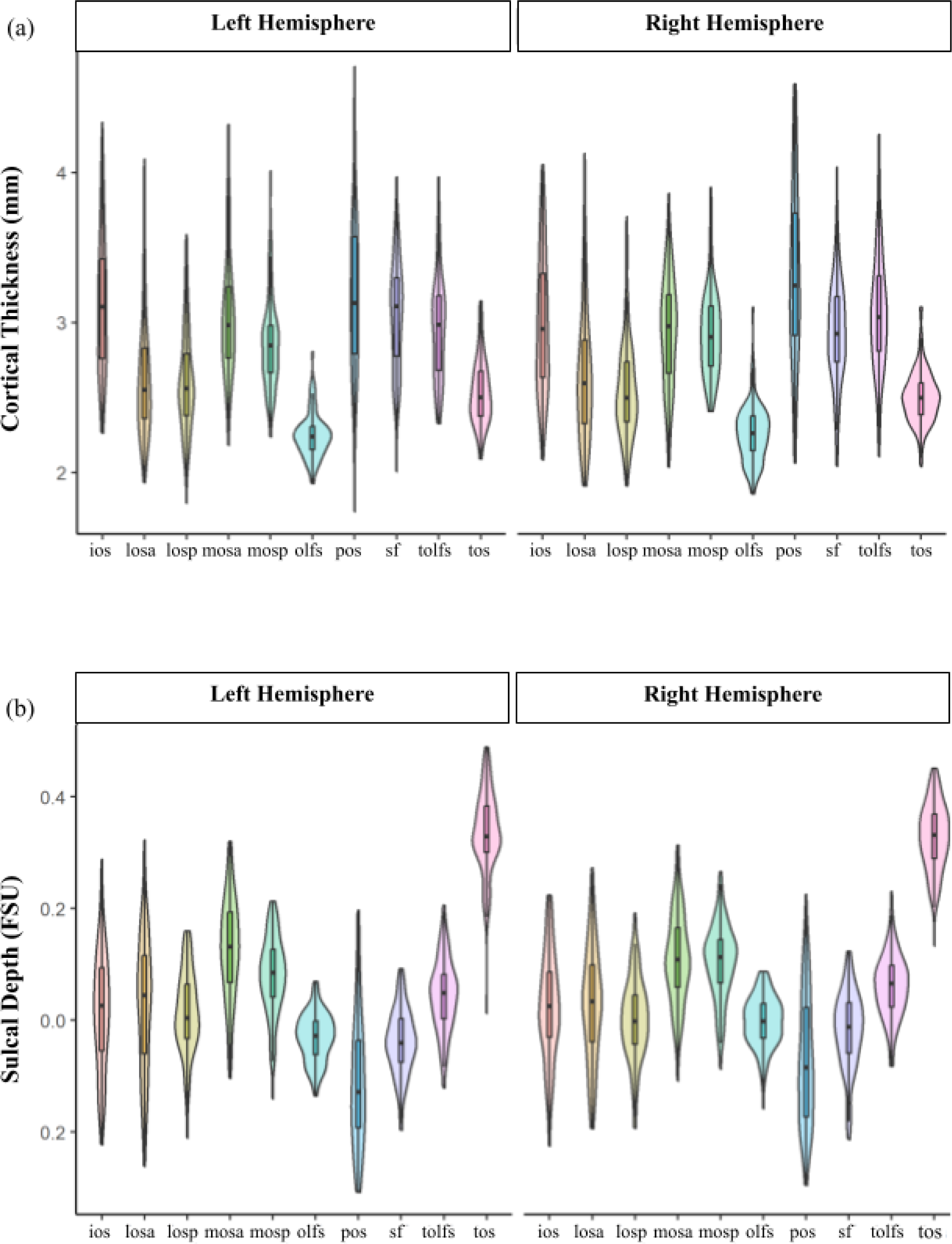
Distributions of OFC sulcal depth and cortical thickness in the present transdiagnostic sample. (a) Violin plots displaying cortical thickness as a function of OFC sulcus (x-axis) and hemisphere (left: left panel, right: right panel). (b) Same as (a), but for sulcal depth.

**Supplementary Figure 3.**
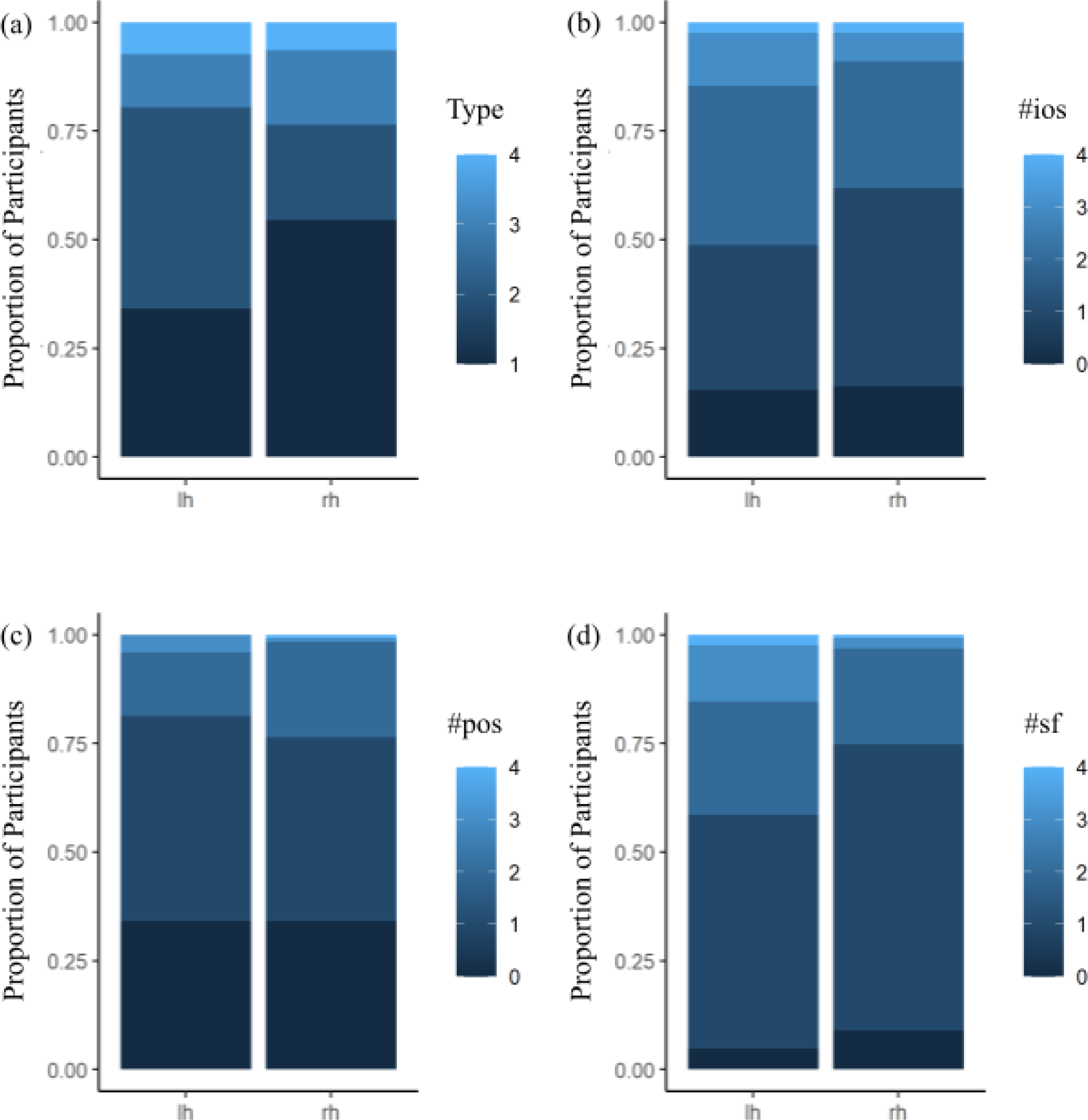
Incidence of sulcogyral type and variable sulcal components in the present transdiagnostic sample. Stacked bars show the proportion of individuals (N = 118) who had each (a) type, (b) number of ios components, (c) number of pos components, (d) number of sf components in the left hemisphere (lh) and right hemisphere (rh). Sulcal abbreviations correspond to those used in Figure 1.

**Supplementary Figure 4.**
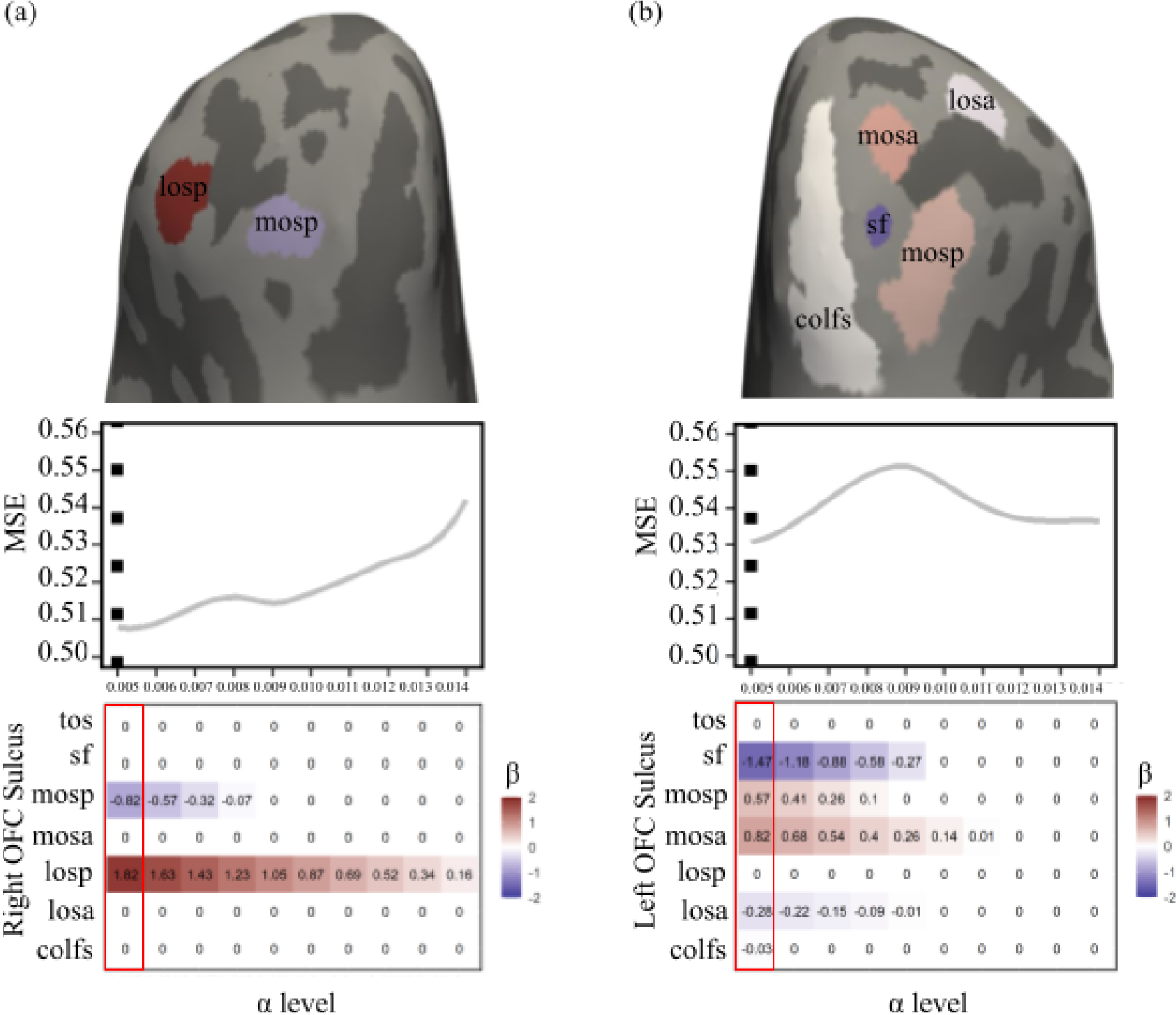
Characteristics of the Olfactory Sulcal Complex (olfs-c) offer little (left hemisphere) to no (right hemisphere) predictive value to the data-driven model. Inflated cortical surfaces including LASSO selected sulci (top) colored according to the beta values (bottom). Line graphs depict MSE at each corresponding alpha value (middle). The dotted line represents the alpha which minimizes MSE. Matrices reflect the beta values of the predictors at each corresponding alpha value (bottom). Beta values used in the model which minimize MSE are outlined in red. Sulcal abbreviations correspond to those used in Figure 1. When including the tolfs and olfs together, the model does not select these sulci in the right hemisphere (a), and the beta value plummets in the left hemisphere (b).

**Supplementary Figure 5.**
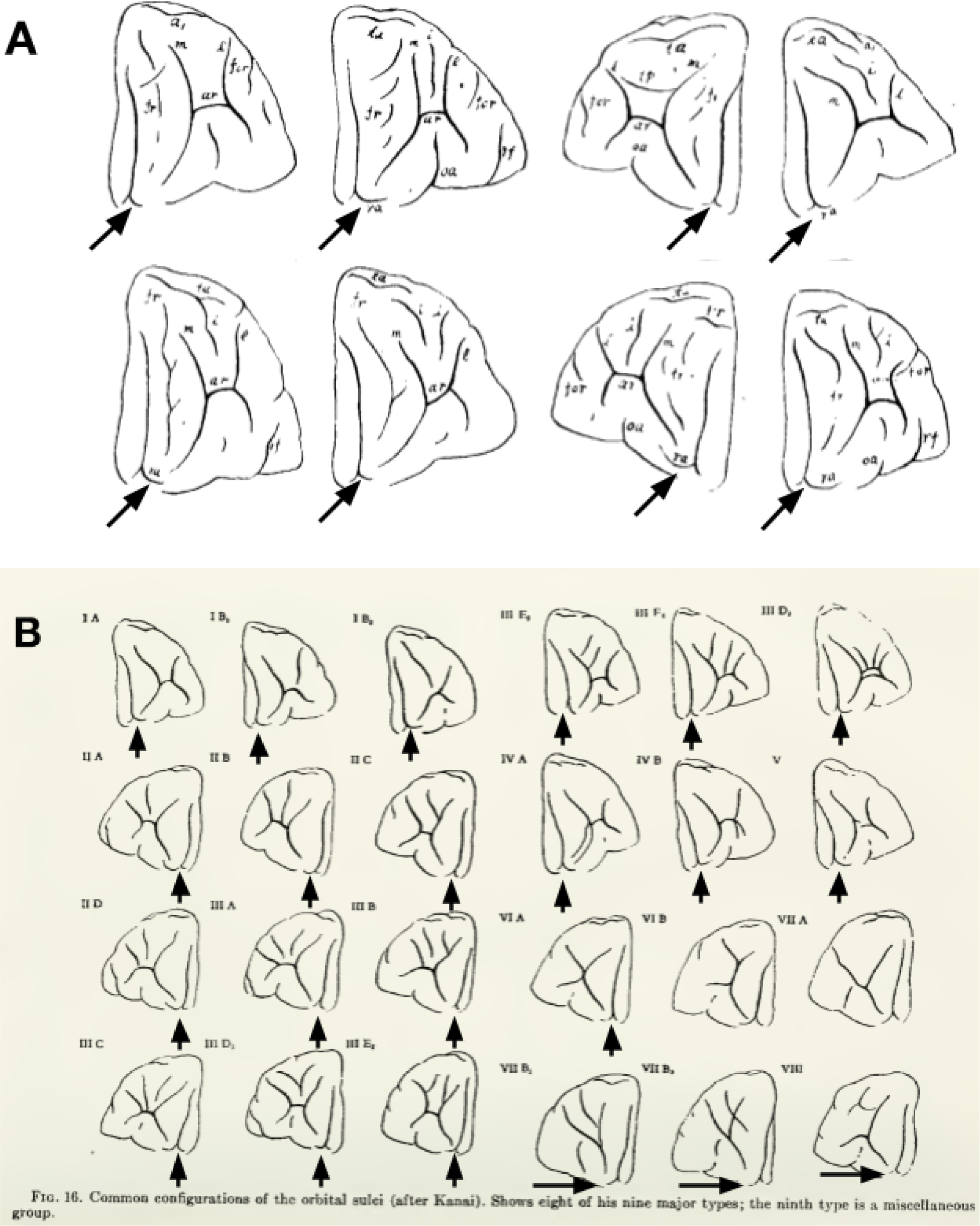
Historical depictions of the transverse olfactory sulcus. (a) Kanai (83) defined a branch, ra, which he referred to as the “lateraler Ast des Sulcus olfactorius,” or lateral branch of the olfactory sulcus (arrow in the image). These are a subset of his images. Note that even when unlabeled, there are clear depictions of medial and lateral rami, which compose our definition of the tolfs. (b) In their *Isocortex of Man,* Bailey and Bonin (84) included the many different tyes of the OFC sulcal patterning documented by Kanai (83). Though unlabeled, the transverse olfs is identifiable in each of their images (black arrows). As documented by Retzius (85), the lateral side is often more prominent than the medial branch.

**Supplementary Figure 6.**
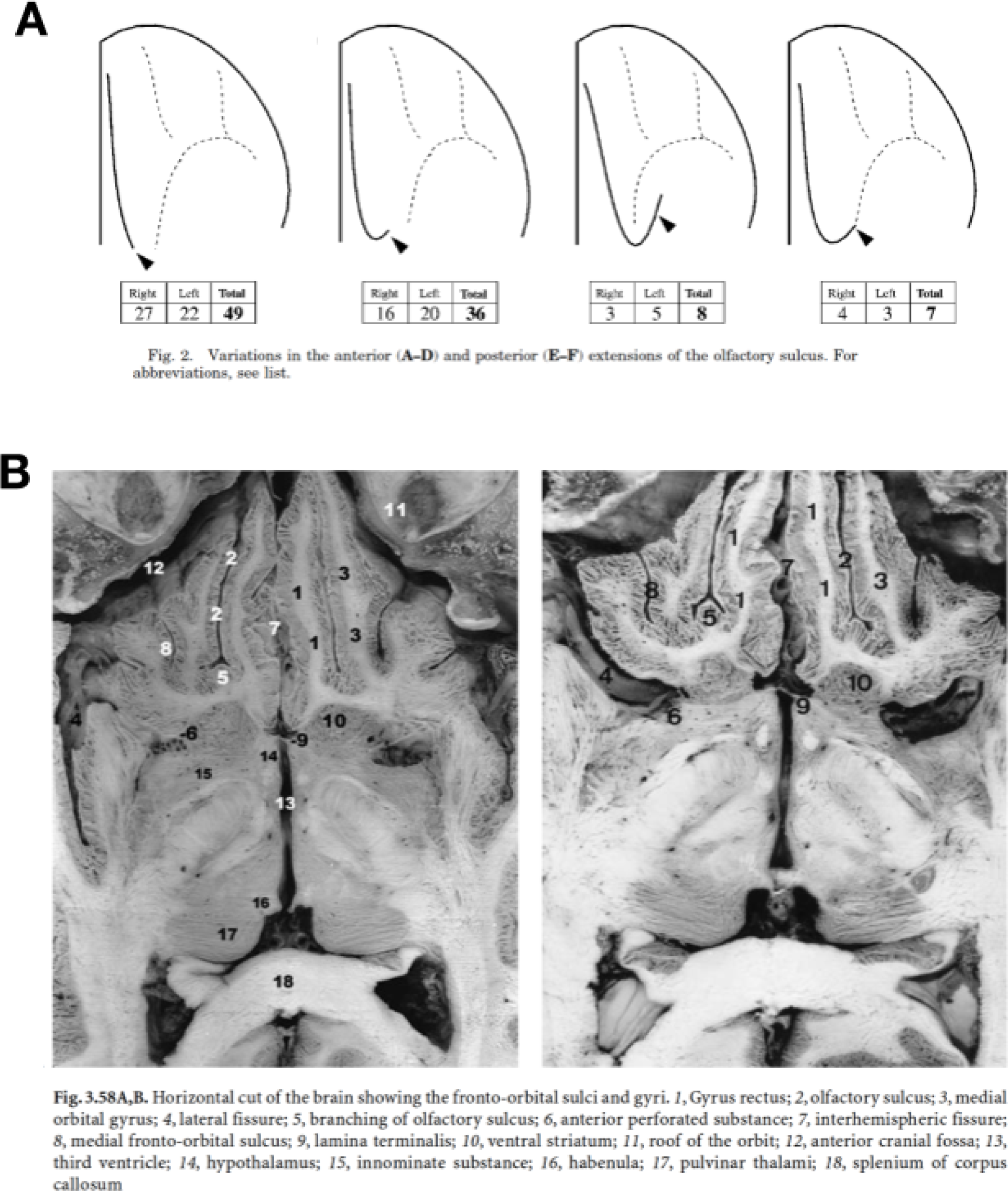
Modern depictions of the transverse olfactory sulcus. (A) Chiavaras and Petrides (34) refer to a posterior hook of the olfactory sulcus (arrowhead in the images). (B) Tamraz and Comair (86) refer to a branching of the olfactory sulcus in the posterior extent (“5” in the included image, with the original caption). Importantly, this also indicates that the tolfs is identifiable in post-mortem brains.

